# Persistent serum protein signatures define an inflammatory subset of long COVID

**DOI:** 10.1101/2022.05.09.491196

**Authors:** Aarthi Talla, Suhas V. Vasaikar, Gregory Lee Szeto, Maria P. Lemos, Julie L. Czartoski, Hugh MacMillan, Zoe Moodie, Kristen W. Cohen, Lamar B. Fleming, Zachary Thomson, Lauren Okada, Lynne A. Becker, Ernest M. Coffey, Stephen C. De Rosa, Evan W. Newell, Peter J. Skene, Xiaojun Li, Thomas F. Bumol, M. Juliana McElrath, Troy R. Torgerson

## Abstract

Long COVID or post-acute sequelae of SARS-CoV-2 (PASC) is a clinical syndrome featuring diverse symptoms that can persist for months after acute SARS-CoV-2 infection. The etiologies are unknown but may include persistent inflammation, unresolved tissue damage, or delayed clearance of viral protein or RNA. Attempts to classify subsets of PASC by symptoms alone have been unsuccessful. To molecularly define PASC, we evaluated the serum proteome in longitudinal samples from 55 PASC individuals with symptoms lasting ≥60 days after onset of acute infection and compared this to symptomatically recovered SARS-CoV-2 infected and uninfected individuals. We identified subsets of PASC with distinct signatures of persistent inflammation. Type II interferon signaling and canonical NF-κB signaling (particularly associated with TNF), were the most differentially enriched pathways. These findings help to resolve the heterogeneity of PASC, identify patients with molecular evidence of persistent inflammation, and highlight dominant pathways that may have diagnostic or therapeutic relevance.

**One Sentence Summary:** Serum proteome profiling identifies subsets of long COVID patients with evidence of persistent inflammation including key immune signaling pathways that may be amenable to therapeutic intervention.

## MAIN TEXT

New, recurrent or prolonged symptoms after acute SARS-CoV-2 infection are termed post-acute sequelae of SARS-CoV-2 (PASC) or long COVID. A recent systematic review of 38 papers reported that one-third or more of surviving COVID-19 patients experienced at least one PASC symptom during the 2-5 months after the onset of acute infection ^1^. PASC symptoms are numerous and varied, impacting virtually every major organ system (https://www.cdc.gov/coronavirus/2019-ncov/long-term-effects/index.html) ^2,3^ and can last for weeks or months. Despite the large number of individuals affected, the lack of consensus diagnostic criteria or standardized outcome measures impede efforts to effectively group persons to establish clinical etiologies or to evaluate outcomes for therapeutic trials ^4^. There are also no clearly defined molecular markers of disease or definitive diagnostic tests. To make matters more complicated, it is recognized that similar clinical symptoms could arise after acute infection regardless of whether they were caused by persistent inflammatory disease initiated by the viral immune response, unresolved organ or tissue damage, or delayed viral clearance. Identification of molecular features capable of mechanistically defining the heterogeneity of PASC could be transformative, allowing clinicians and researchers to better subset patients and highlighting potential targets for therapeutic intervention.

We hypothesized that the serum proteome may provide insights into potential drivers of PASC symptomatology and may offer a clinically accessible tool to help define subgroups of PASC. We therefore analyzed the serum proteome using the Olink Explore 1536 panel in 55 adults (21 men, 34 women; age 22-82 years) with persistent symptoms lasting ≥60 days after an acute, PCR-confirmed SARS-CoV-2 infection (termed “PASC”), in 24 (9 men, 15 women; age 20-79 years) who symptomatically recovered after a PCR-confirmed SARS-CoV-2 infection (termed “Recovered”), and in 22 (12 men, 10 women; age 29-77) who had a negative nasopharyngeal PCR test (termed “Uninfected”) (**Fig S1A**). The uninfected individuals had blood drawn once at study entry while the PASC and recovered persons had one or more blood draws at timepoints ≥60 days and up to 379 days post-symptom onset (PSO) of acute COVID (**Fig S1B**). Most patients had mild symptoms during acute infection (World Health Organization (WHO) ordinal scale 2 or 3) but 3 subjects with moderate scores were hospitalized and required oxygen (WHO ordinal scale 5) ^5^. None required mechanical ventilation. All were unvaccinated. Symptoms of each PASC patient are provided in **Table S1**.

Previous studies have divided PASC patients into subsets based on either type, number, or severity of clinical features ^6,7^. For our cohort, hierarchical clustering on PASC symptomatology alone at ≥60 days PSO did not clearly drive significant patient clustering (**Fig S1A, S2A**). We next attempted to use symptoms to drive clustering of significantly associated serum protein signatures, but no single symptom or combination of symptoms was able to clearly distinguish patient groups (**Fig S2B, C, D**) suggesting that symptoms alone are unable to differentiate subsets of PASC.

We therefore took an alternative approach, using unbiased clustering of the serum proteome across the entire cohort (PASC + recovered + uninfected) to find clusters of individuals that had similar serum proteome signatures regardless of their COVID status or symptomatology.

Canonical pathway enrichment was performed on the first post-60 day sample available for each PASC subject, the last available post-60 day sample for each recovered subject (to maximize the chance that proteome alterations had returned to baseline) and on the solitary sample from the uninfected individuals (see Methods for details). We used the curated canonical pathways from the Molecular Signatures Database (MSigDB) and applied a rule-in approach ^8^, which resulted in 85 pathways that distinguished PASC from recovered and uninfected individuals with a significant rule-in performance (p < 0.01). These pathways were merged into 54 modules to avoid gene set redundancy using the enrichment map approach with a minimum Jaccard index threshold of 25% (**Table S3**) (see Methods). Hierarchical clustering using the 54 proteomic modules identified 5 discrete clusters that showed distinct expression patterns of the modules (**Fig 1A**). Two of the clusters (4 & 5) showed a marked enrichment for inflammatory modules while clusters 1, 2, and 3 lacked a distinct inflammatory protein signature. Inflammatory clusters 4 and 5 included predominantly PASC individuals (91% and 80% respectively) whereas cluster 1 consisted of only uninfected or recovered individuals. Clusters 2 and 3 consisted of a mixture of PASC (48% and 28% respectively), recovered, and uninfected subjects (**Fig S3A**). The distribution of PASC subjects across inflammatory (4 & 5; 65% of PASC) and non-inflammatory (2 & 3; 35% of PASC) proteomic clusters underscores the heterogeneity of PASC. To determine whether the differential serum proteomic signatures discovered by comparing the first post-60 day PSO sample for PASC to the last post-60 day PSO sample for recovered are stable over time, we extended our analysis to include all longitudinal samples available for each subject. We found that PASC subjects exhibiting an inflammatory protein signature continue to have that signature over time and that most subjects remained in the same cluster throughout the study period (**Fig S3B**). Whether the two clusters of inflammatory PASC described here represent two distinct subtypes with different molecular drivers or a continuum of disease requires testing in future studies.

**Figure 1:**
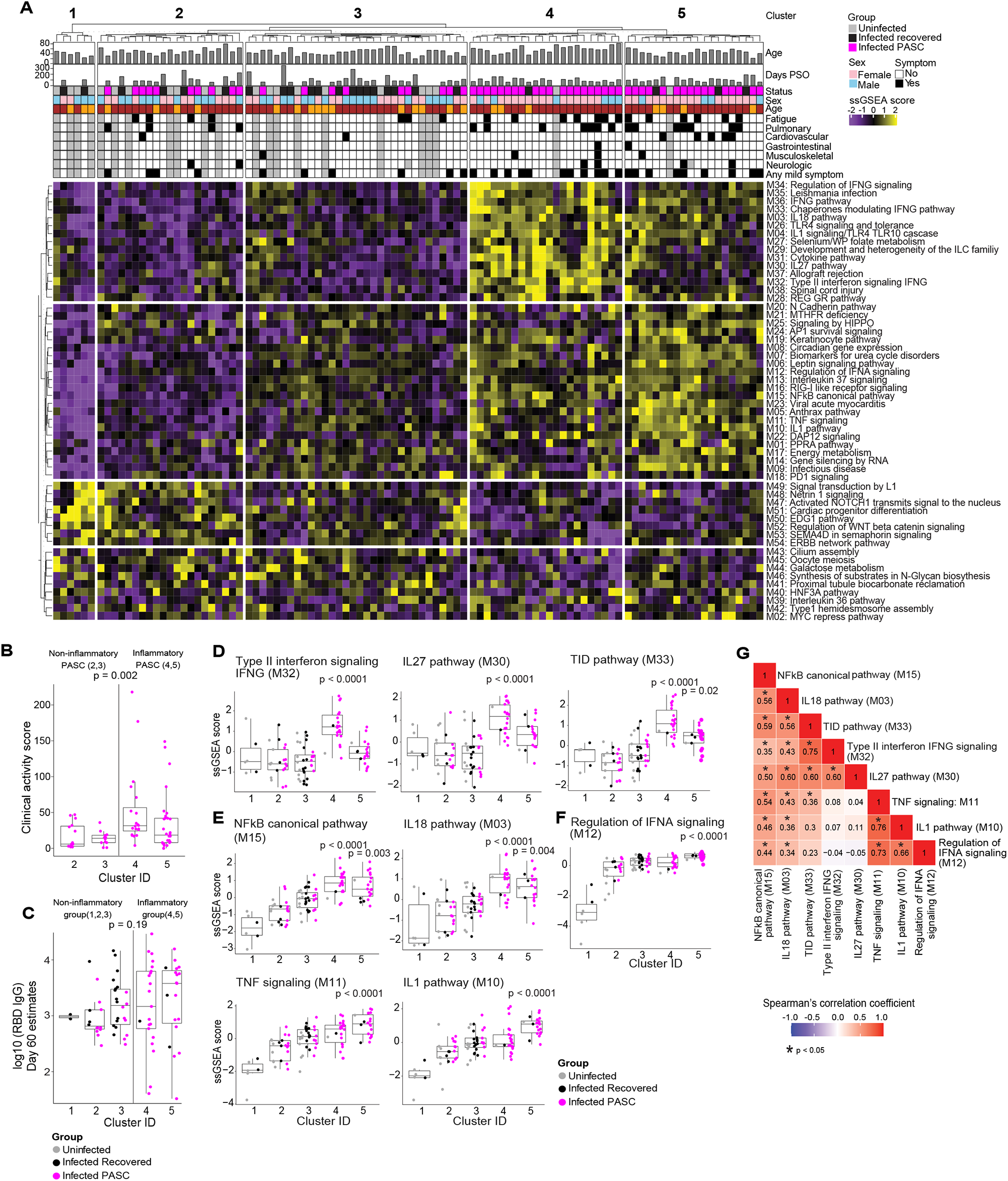
Serum proteomic clustering of PASC. (A) Heatmap of the rule-in method based unsupervised clustering of Olink serum proteome data across all patients in the cohort (PASC + recovered + uninfected). Rows represent modules, columns represent samples and the scaled ssGSEA module score across samples is depicted from low (purple) to high (yellow). The method identifies 2 clusters of subjects with higher inflammatory module signatures (4 & 5) relative to the other three clusters of subjects (1, 2, 3) that lack inflammatory signatures.

We hypothesized that an inflammatory plasma protein signature may also correlate with being more symptomatic during acute infection. However, because this study cohort primarily experienced mild acute COVID-19 symptoms (WHO ordinal scale 2 or 3), commonly used COVID severity indices did not capture a range of heterogeneity in symptomatology. We therefore developed a clinical activity score for the acute phase of mild COVID that accounted for both duration of symptoms and their impact on activities of daily (see Methods).

Inflammatory PASC subjects in clusters 4 & 5 had a significantly higher pre-PASC clinical activity score (Wilcoxon rank sum p=0.002) compared to non-inflammatory PASC subjects in clusters 2 & 3 (**Fig 1B**). We wondered whether subjects with an inflammatory protein signature may have mounted less robust immune responses to SARS-CoV-2, thus potentially delaying viral clearance or increasing risk for viral persistence. However, comparison of SARS-CoV-2 receptor binding domain (RBD)-specific IgG titers in infected subjects (PASC + Recovered) 60 days PSO identified no significant difference between the inflammatory (4 and 5) and non-inflammatory (1, 2 and 3) clusters (**Fig 1C**). Comparison of SARS-CoV-2-specific CD4+ and CD8+ T cell frequencies ^9,10^ between the inflammatory and non-inflammatory PASC also did not show any significant difference (**Fig S3C, Fig S3D**).

Among the 54 modules that defined the 5 clusters (**Fig 1A**), we identified those that significantly distinguished each cluster by calculating the single-sample Gene Set Enrichment Analysis (ssGSEA) score per module across samples (**Table S4**). Ranking modules by adjusted p-value identified those most significantly associated with clusters 4 and 5 (**Table S5, Fig S4, Fig S5**).

Within cluster 4, multiple pathways associated with type II interferon (IFN-γ) signaling (Type II IFN signaling, IL-27, TID, etc.) were among those most highly enriched (**Fig 1D**). Canonical NF-κB signaling and NF-κB activating cytokine pathways (IL-18, TNF, IL-1 were enriched in both clusters 4 and 5 (**Fig 1E**). In addition, cluster 5 was also enriched for proteins associated with regulation of IFN-α signaling (**Fig 1F**). The expression scores of these modules across all samples were significantly correlated with each other, indicating, patients with higher IFN-γ signaling have higher IL27, IL18, and NF-κB signaling, and patients with higher TNF signaling have higher IL1, NF-κB, and IFN-α signaling, suggesting a global activation of immune cascades that drive inflammation (**Fig 1G**).

We next investigated the individual proteins differentially expressed in the serum of subjects within each cluster. Clusters 1-5 were individually compared to all other clusters. Cluster 4 had 234 differentially expressed proteins (DEPs) whereas cluster 5 had 296 DEPs (**Table S6**; adj. p-value <0.05). Since cytokines, chemokines, and cytokine/chemokine receptors are major drivers of inflammation and potential targets for therapeutic intervention, we focused on these and ranked individual DEPs by adjusted p-value (**Fig 2A**). IFN-γ was found to be the cytokine that most significantly defines cluster 4. Moreover, IFN-γ was the top DEP enriched in cluster 4 among all 1463 analytes in the Olink protein panel (**Fig S6, S7, Table S6**). Increased expression of chemokines and cytokines known to be regulated by IFN-γ including CXCL9, CXCL10, CXCL11, and IL-27 in cluster 4 suggests that it is functionally active (**Fig 2A, 2B**). We also observed increased expression of IL-12 p40 (IL12B) and the IL-12 p40/p70 heterodimer (IL12A_IL12B) in cluster 4, which may drive expression of IFN-γ and an overall Th1 signature.

**Figure 2:**
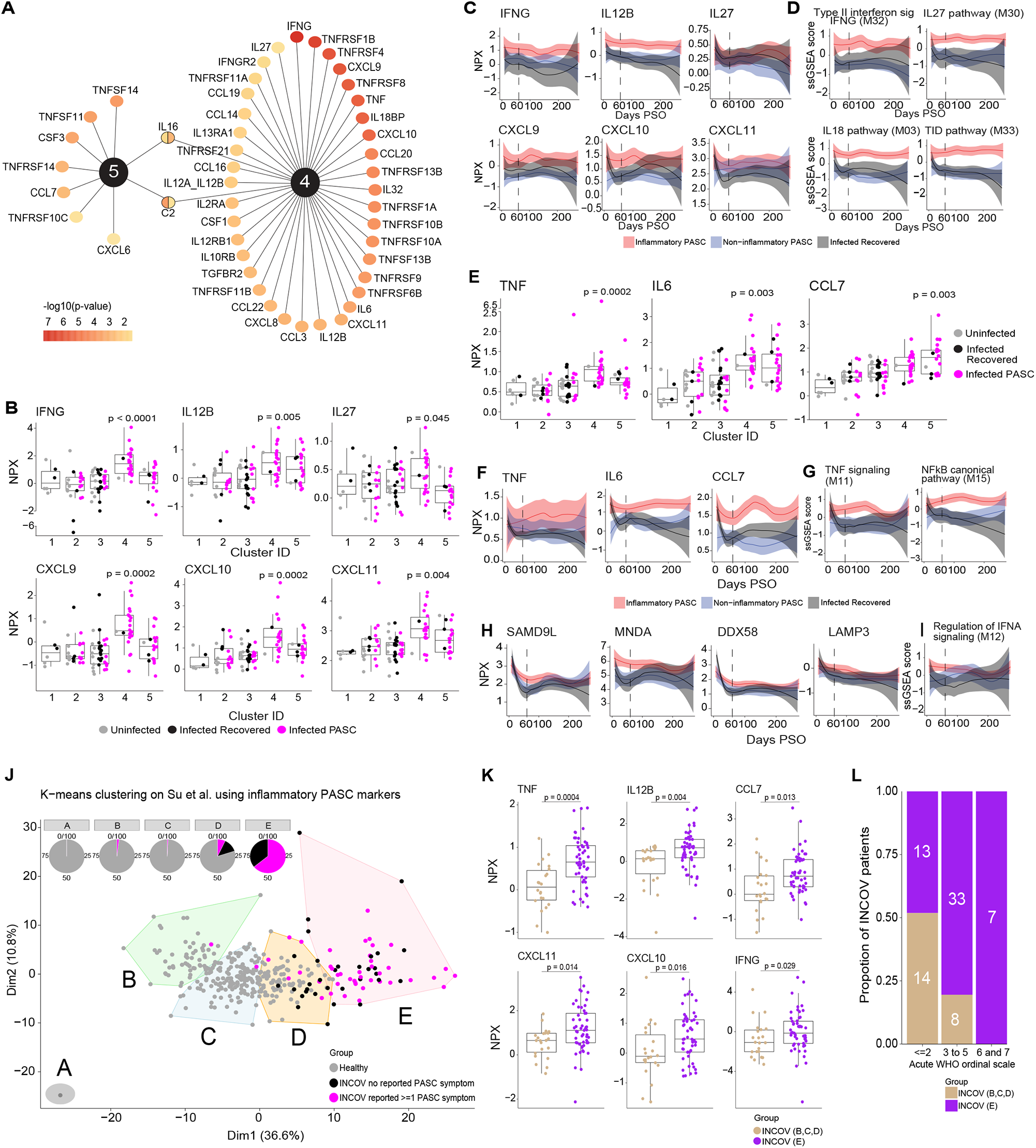
Key protein signals driving inflammatory PASC signatures. (A) Top ranked differentially expressed cytokines, chemokines, and cytokine/chemokine receptors by adjusted p-value of <0.05 that are associated with inflammatory protein clusters 4 & 5. The color gradient of each node represents the -log10 adjusted p-value. (B) Box and jitter plots of olink Normalized Protein Expression (NPX) (y-axis) of IFN-γ and its related cytokines and chemokines across clusters (x-axis) that were significantly upregulated exclusively in cluster 4. P-values determined by Wilcoxon rank sum test were calculated comparing inflammatory cluster 4 and inflammatory cluster 5 independently to clusters 1,2,3. (C) Longitudinal loess fit plots of Olink NPX of IFN-γ and its related cytokines and chemokines on samples available from early acute infection through >60 days PSO (x-axis). PASC patients from the inflammatory clusters 4 and 5 are represented here as inflammatory PASC (red), PASC patients from clusters 2 and 3 are represented here as non-inflammatory PASC (blue) while the recovered patients are represented in black. (D) Longitudinal loess fit plots of the ssGSEA scores (y-axis) of IFN-γ related modules over time (x-axis). (E) Box and jitter plots of Olink NPX (y-axis) expression levels of TNF, IL6 and CCL7 across clusters (x-axis) that were significantly differentially upregulated clusters 4 and 5. P-values determined by Wilcoxon rank sum test were calculated comparing inflammatory cluster 4 and inflammatory cluster 5 independently to clusters 1,2,3. (F) Longitudinal loess fit plots of Olink NPX (y-axis) of TNF, IL6 and CCL7 over time (x-axis). (G) Longitudinal loess fit plots of the ssGSEA scores (y-axis) of TNF and NF-κB related signaling modules over time (x-axis). (H, I) Longitudinal loess fit plots of Olink NPX and ssGSEA scores (y-axes) of type-I IFN-driven proteins and the IFN-α module overtime (x-axis) respectively. (J) K-means unsupervised clustering of Olink proteomic data from Su Y et al (2022) showing 5 clusters of INCOV patients and healthy controls. Pie charts show the percentage of each cluster consisting of INCOV patients and healthy subjects. (K) Cytokines/chemokines significantly upregulated in the INCOV cluster E vs. INCOV from clusters B,C, and D. P-values were determined by a Wilcoxon rank sum test. (L) Distribution of different disease severities (as judged by WHO ordinal scale) across INCOV patients in cluster E vs INCOV patients in clusters B,C,D. Y-axis and the numbers in bar graphs represent proportion and number of patients per INCOV group in each WHO scale bin respectively.

To determine whether IFN-γ and IFN-γ driven cytokines, chemokines, and pathways remained persistently elevated over time in inflammatory PASC, we evaluated these signatures longitudinally in available samples beginning from early acute infection to 275 days PSO. IFN-γ, IL-12 p40, and IFN-γ-driven chemokines were consistently elevated within inflammatory PASC from clusters 4 & 5 compared to non-inflammatory PASC from clusters 1, 2, and 3, extending to at least 275 days after initial SARS-CoV-2 infection (**Fig 2C, Fig S8**). IFN-γ related signaling modules also showed persistent enrichment over the same time (**Fig 2D, Fig S9**).

In addition to IFN-γ, TNF, TNF-driven cytokines and chemokines (including IL-6 and CCL7 (MCP3)), and several TNF receptor superfamily members were also increased in clusters 4 and 5 (**Fig 2A, 2E, Fig S8**). TNF, IL-6, and CCL7 remained persistently elevated in inflammatory PASC over time compared to non-inflammatory PASC (**Fig 2F, Fig S8**). In addition, TNF signaling and canonical NF-κB signaling pathways previously found to be enriched at early time points in inflammatory PASC remained elevated over time (**Fig 2G, Fig S9**).

Finally, the pathway related to expression of IFNA signaling was found to be enriched at the first post-60 day PSO timepoint in cluster 5 (**Fig 1F**). The Olink assay only quantifies IFN-γ and IFNλ1 but we observed increased expression of proteins associated with type I IFN activation including SAMD9L, MNDA, DDX58, LAMP3, and others (**Fig S6, Fig S7**). These proteins were found to be highly increased early after acute infection but in inflammatory PASC, remained elevated over time compared to non-inflammatory PASC. Longitudinal assessment showed that SAMD9L, MNDA, DDX58, and LAMP3 trended toward the levels seen in non-inflammatory PASC and recovered subjects by approximately 180 days post infection (**Fig 2H**), similar to the kinetics observed for the expression of IFNA signaling pathway over time (**Fig 2I**). This is notable in light of recent studies reporting detection of SARS-CoV-2 RNA and protein in gastrointestinal and hepatic tissue of convalescent patients up to 180 days after acute infection and in diverse extrapulmonary tissues including brain up to 230 days after acute symptom onset ^11,12^. Whether residual viral RNA and/or protein may serve as a driver of the phenotype in inflammatory PASC remains to be investigated more thoroughly.

To determine whether our observations could be extended to an independent cohort of PASC patients collected across a broader range of acute COVID severities, we applied a similar analysis approach to the recently published INCOV cohort that included Olink plasma proteomic data from 204 SARS-CoV-2-infected patients and 289 healthy controls ^13,14^. Of the 204 INCOV patients, 75 met the criteria used for our cohort (Olink data available from sample obtained ≥60 days after acute infection + clinical data available). Forty-three (57%) of these had 1 or more PASC symptoms like the PASC subjects in our cohort and the remainder had no recorded PASC symptoms, similar to the “recovered” group in our cohort. The Olink panel employed in the INCOV study measured only 443 of the 1472 proteins measured in our study but 163 proteins overlapped with the inflammatory signatures that significantly defined clusters 4 & 5 in our cohort. To be consistent with our cohort, k-means unsupervised clustering of the Olink proteomic data from the INCOV cohort was performed with k=5 using the 163 overlapping proteins on the sample available at the first timepoint ≥60 days PSO per INCOV patient (n=75).

Of the 5 INCOV clusters, cluster E is similar to our inflammatory PASC clusters 4 and 5 with significant enrichment of 129 of the 163 proteins (79%) that defined our inflammatory PASC. Similar to our clusters 4 and 5 that consisted primarily of PASC persons, 64.2% of the persons in INCOV cluster E were patients with persistent PASC symptoms (**Fig 2J, Fig S10, Table S7**). No healthy controls clustered with cluster E. INCOV cluster D was made up of a mixture of PASC, recovered, and healthy controls, like our cluster 2. The remaining clusters (A, B, C) were made up predominantly of healthy individuals. Among the cytokines and chemokines observed in our inflammatory PASC, proteins that were also significantly higher in INCOV cluster E included IFN-γ, IL12, CXCL10, CXCL11, TNF, and CCL7 (**Fig 2K**) that were increased in our cluster 4 along with others increased in our cluster 5 (DDX58, LAMP3, etc,) (**Fig S10, Table S7**). Lastly, the broader diversity of disease severity in the INCOV cohort compared to our mild to moderate cohort, allowed us to make an association between the clinical measure of acute disease severity (WHO ordinal scale score) and proteomic inflammatory signatures. Interestingly, INCOV patients from cluster E predominantly exhibited an acute WHO ordinal score of ≥3 reflecting the association between more severe acute disease and persistent inflammation ^15^ (**Fig 2L**).

These findings substantially extend previous observations that have variably reported increased expression of IFN-γ, IFN-β, IFN-λ1/2/3, TNF, IL-6, IL-1β, and PTX3 in plasma from PASC patients using targeted cytokine panels ^16–18^. While previous studies grouped all PASC participants, we provide the first evidence that more than half of all PASC have an inflammatory protein signature while others do not have this signature. We show that in inflammatory PASC, the IL-12/IFN-γ axis is highly active and is combined with a NF-κB driven protein signature, possibly driven by TNF and leading to excess IL-6 expression. Furthermore, we show evidence of a persistent type I IFN driven protein signature present in inflammatory PASC group that trends toward normal approximately 6 months post-infection, paralleling recent reports of persistent SARS-CoV-2 RNA and protein being detected in non-pulmonary tissues up to 6-8 months after infection ^11,12^. We show that these findings can be applied to another PASC proteomic dataset to identify PASC subjects with persistent inflammatory disease. These data suggest a serum protein signature that could be used diagnostically to address the challenge of clinical heterogeneity in PASC. It also provides insights to potential molecular mechanisms of disease and possible therapeutic targets within individuals with an inflammatory protein signature (TNF, IL-6, IFN-γ, etc.).

Metadata including age, sex, symptoms, days post-symptom onset (PSO) are shown at the top of the heatmap. (B) Clinical activity score of mild acute COVID symptoms in PASC subjects from inflammatory (4 & 5) vs. non-inflammatory (2 & 3) clusters. The p-value determined by Wilcoxon rank sum test was calculated comparing, as a group, inflammatory PASC vs non-inflammatory PASC. (C) Receptor binding domain (RBD)-specific IgG titers in PASC and recovered patients within each cluster at 60 days PSO. The p-value determined by Wilcoxon rank sum test was calculated comparing, as a group, inflammatory clusters vs non-inflammatory clusters. (D-F) Box and jitter plots of the ssGSEA scores (y-axis) across all clusters (x-axis) for the top ranked modules that were enriched in inflammatory clusters 4 and 5. P-values determined by Wilcoxon rank sum test were calculated comparing inflammatory cluster 4 and inflammatory cluster 5 independently to clusters 1,2,3. (G) Pair-wise Spearman’s correlation coefficient heatmap between top enriched modules that define inflammatory clusters 4 and 5 demonstrating co-enrichment of modules.

## MATERIALS AND METHODS

### Regulatory approvals

COVID19 Fred Hutch samples and healthy controls: FH RG: 1007696 IR File: 10440 Main Consent 04/05/2020 and 6/04/2020 Seattle COVID-19 Cohort Study to Evaluate Immune Responses in Persons at Risk and with SARSCoV-2 Infection.

### Study Conduct

Serum was collected from participants enrolled in the longitudinal study, “Seattle COVID-19 Cohort Study to Evaluate Immune Responses in Persons at Risk and with SARS-CoV-2 Infection”. Eligibility criteria included adults in the greater Seattle area at risk for SARS-CoV2 infection or those diagnosed with SARS-CoV-2 by a commercially available SARS CoV-2 PCR assay. Study data were collected and managed using REDCap electronic data capture tools hosted at Fred Hutchinson Cancer Research Center, including detailed information on symptoms during acute infection and longitudinal follow-up ranging from 33-379 days post symptom onset. All but 2 persons in the “uninfected” group had at least 1 symptom of SARS-CoV-2 infection within the 14 days prior to study screening but had a negative SARS-CoV-2 nasopharyngeal PCR test. Informed consent was obtained from all participants at the Seattle Vaccine Trials Unit and the Fred Hutchinson Cancer Research Center Institutional Review Board approved the studies and procedures.

### Symptoms category clustering

We collected symptom information from each donor over multiple visits. The symptom data were merged into six major categories including fatigue/malaise, pulmonary, cardiovascular, gastrointestinal, musculoskeletal, and neurologic. Other mild symptoms were combined into a single category as “any mild symptoms” (Table S1). Symptom information was converted to binary format where yes=1 and no=0. Missing symptom information is denoted by NA. The binary information was used to perform principal component analysis (PCA) and visualize sample clustering using factoextra (v1.0.7). The contribution of variation for each symptom category was retrieved and shown in bar plot. For each symptom category we identified symptom-specific differential plasma proteins using linear mixed model ^19^. We used lme4 package (v1.1) to carry out linear mixed model analysis where age, sex were fixed variables and donor information was a random variable.

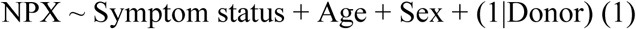

The p-value is obtained from chi-square statistics. The specific symptom category associated with differential plasma proteins selected using p < 0.05. The identified differential proteins from six symptom specific categories were merged together and their expression visualized in a heatmap using package ComplexHeatmap (v2.4).

### Symptom activity metrics and scoring for mild to moderate acute COVID symptoms

Symptom activity in mild to moderate acute COVID was classified by participant report of impact on Activities of Daily Living (ADLs) for each day of illness (Health et al. 2017). Days hospitalized were recorded as were any treatment or therapies received. Participants were scored according to their maximum symptom activity for each day: 0, no symptoms; 1, mild impact on ADLs reported; 2, moderate impact on ADLs reported; 3, severe illness without hospitalization; 4, severe illness with hospitalization; 5, hospitalized with ICU care, or 6, life threatening illness. Duration was assigned for days spent at each level of symptom activity. A clinical activity score was calculated for each subject by multiplying the symptom activity score by the number of days spent at each level, then summing all values.

### Sample processing

Blood was drawn into a serum separator tube and serum samples were processed, aliquoted and frozen within 4 hours of blood draw.

### Olink serum protein measurement

Serum samples were inactivated with 1% Triton X-100 for 2h at room temperature according to the Olink COVID-19 inactivation protocol. Inactivated samples were then run on the Olink Explore 1536 platform, which uses paired antibody proximity extension assays (PEA) and a next generation sequencing (NGS) readout to measure the relative expression of 1472 protein analytes per sample. Analytes from the inflammation, oncology, cardiometabolic, and neurology panels were measured.

For plate setup, samples were randomized across plates to achieve a balanced distribution of age and gender. Longitudinal samples from the same participant were run on the same plate. To facilitate comparisons with future batches, sera from 15 donors was commercially purchased (BioIVT) and randomly interspersed amongst the above study samples. Commercial samples included serum from COVID-19 serology-negative, serology-positive, PCR-positive, and recovered (no longer symptomatic) participants.

Data were first normalized to an extension control that was included in each sample well. Plates were then standardized by normalizing to inter-plate controls run in triplicate on each plate. Data were then intensity normalized across all samples. Final normalized relative protein quantities were reported as log2 normalized protein expression (NPX) values.

### Olink data preprocessing

Olink results and QC flags were reviewed for overall quality. Results for TNF, IL6 and CXCL8, which were measured on all 4 Olink panels, were reviewed prior to averaging to a single NPX value for analysis. Two samples had discrepant cross-panel measurements on these proteins. The results that trended most consistently with the participant’s longitudinal measurements were kept and averaged. Serum samples were analyzed in two batches. Following the method recommended by Olink, results of the later batch were bridged to those of the earlier batch using a set of 42 cohort samples that were tested in both batches. A batch offset for each analyte was calculated as the median difference on the 42 samples as measured between the two batches, excluding samples with QC warning flags. The analyte-specific offsets were then added to the raw NPX values of the later batch.

### Antibody ELISAs for RBD

Half-well area plates (Greiner) were coated with purified RBD protein at 16.25ng/well in PBS (Gibco) for 14-24h at room temperature. After 4 150ul washes with 1X PBS, 0.02% Tween-2 (Sigma) using the BioTek ELx405 plate washer, the IgA and IgG plates were blocked at 37°C for 1-2 hours with 1X PBS, 10% non-fat milk (Lab Scientific), 0.02% Tween-20 (Sigma); IgM plates were blocked with 1X PBS, 10% non-fat milk, 0.05% Tween-20. Serum samples were heat inactivated by incubating at 56°C for 30 minutes, then centrifuged at 10,000 x g / 5 minutes, and stored at 4°C previous to use in the assay. For IgG ELISAs, serum was diluted into blocking buffer in 7-12 1:4 serial dilutions starting at 1:50. For IgM and IgA ELISAs, serum was diluted into 7 1:4 serial dilutions starting at 1:12.5 to account for their lower concentration. A qualified pre-pandemic sample (negative control) and a standardized mix of seropositive serums (positive control) was run in each plate and using to define passing criteria for each plate. All controls and test serums at multiple dilutions were plated in duplicate and incubated at 37°C for 1 hour, followed by 4 washes in the automated washer. 8 wells in each plate did not receive any serum and served as blocking controls. Plates then were plated with secondary antibodies (all from Jackson ImmunoResearch) diluted in blocking buffer for 1h at 37C. IgG plates used donkey anti-human IgG HRP diluted at 1:7500; IgM plates used goat anti-human IgM HRP diluted at 1:10,000; IgA plates used goat anti-human IgA HRP at 1:5000. After 4 washes, plates were developed with 25ul of SureBlock Reserve TMB Microwell Peroxide Substrate (Seracare) for 4 min, and the reaction stopped by the addition of 50ml 1N sulfuric acid (Fisher) to all wells.

Plates were read at OD_450nm_ on SpectraMax i3X ELISA plate reader within 20 min of adding the stop solution. OD_450nm_ measurements for each dilution of each sample were used to extrapolate RBD endpoint titers when CVs were less than 20%. Using Excel, endpoint titers were determined by calculating the point in the curve at which the dilution of the sample surpassed that of 5 times the average OD_450nm_ of blocking controls + 1 standard deviation of blocking controls. RBD titers at day 60 PSO were estimated by a linear mixed effects model of titers over time from day 30 PSO with random effects for the intercept and slope, using lme from the nlme R package.

### Intracellular Cytokine Staining (ICS) Assay

Flow cytometry was used to examine SARS-CoV-2-specific CD4+ and CD8+ T-cell responses using a validated ICS assay. The assay was similar to a published report ^20,21^. Peptide pools covering the structural proteins of SARS-CoV-2 were used for the six-hour stimulation. Peptides matching the SARS-CoV-2 spike sequence (316 peptides, plus 4 peptides covering the G614 variant) were synthesized as 15 amino acids long with 11 amino acids overlap and pooled in 2 pools (S1 and S2) for testing (BioSynthesis). All other peptides were 13 amino acids overlapping by 11 amino acids and were synthesized by GenScript. The peptides covering the envelope (E), membrane (M) and nucleocapsid (N) were initially combined into one peptide pool, but the majority of the assays were performed using a separate pool for N and one that combined only E and M. Several of the open reading frame (ORF) peptides were combined into two pools, ORF 3a and 6, and ORF 7a, 7b and 8. All peptide pools were used at a final concentration of 1 microgram/ml for each peptide. As a negative control, cells were not stimulated, only the peptide diluent (DMSO) was included. As a positive control, cells were stimulated with a polyclonal stimulant, staphylococcal enterotoxin B (SEB). Cells expressing IFNγ and/or IL-2 and/or CD154 were the primary immunogenicity endpoint for CD4+ T cells and cells expressing IFNγ were the primary immunogenicity endpoint for CD8+ T cells. The overall response to SARS-CoV-2 was defined as the sum of the background-subtracted responses to each of the individual pools. A sample was considered positive for CD4+ or CD8+ T cell responses to SARS-CoV-2 if any of the CD4+ or CD8+ T cell responses to the individual peptide pool stimulations was positive.

Positive responses to a given peptide pool stimulation were determined using the MIMOSA (Mixture Models for Single-Cell Assays) method (Finak et al., 2014a). The MIMOSA method uses Bayesian hierarchical mixture models that incorporate information on cell count and cell proportion to define a positive response by comparing peptide-stimulated cells and unstimulated negative controls. MIMOSA estimates the probabilities that peptide-stimulated responses are responders and applies a false-discovery rate multiplicity adjustment procedure (Newton et al 2004). Responses with false-discovery rate q-values < 0.05 were considered positive. The total number of CD4+ T cells must have exceeded 10,000 and the total number of CD8+ T cells must have exceeded 5,000 for the assay data to be included in the analysis.

### Identification of pathways with high rule-in performance

Partial area under the receiver operating characteristic curve (pAUC) ^22,23^. was used to evaluate the rule-in performance ^8^ of individual pathways in identifying PASC subjects with respect to recovered and uninfected subjects. The pAUC bounded by a specificity between 90-100% and the corresponding 99% confidence interval (two-sided) of each pathway were calculated using the “*ci*.*auc*” function in the R package *pROC* with the following parameters: partial.auc=c(0.9, 1), conf.level=0.99, boot.n=1000. A pathway was identified as significant with p < 0.01 if its pAUC lower confidence bound was above the corresponding pAUC of a random, non-performing classifier, i.e. 0.005.

We collected the canonical pathway “c2.cp.v7.2.symbols” geneset and associated gene information from MsigDB (v7.2). The canonical pathway consists of 2871 pathways used to perform single sample GSEA (ssGSEA) using GSVA (v1.40) R package (Hänzelmann et al., 2013 PMID:23323831). Among 2871 pathways, 1960 pathways with overlapping plasma proteins were used as input for GSVA with min.size 2 and max.size 2000 genes as parameters. The ssGSEA resulted in a normalized enrichment score (NES) for each pathway. One sample for each PASC donor was selected as the last time point of infected recovered with >60 days PSO (n=24) and first time point with >60 days PSO for infected PASC donors (n=55). Total 101 donors with one sample including uninfected (n=22) were considered for biomarker analysis.

Rule-in approach implemented to identify pathways significantly associated with PASC donors. Parameters such as confidence interval (CI), pAUC and bootstrap (boot.n) of 200 were used.

Bootsrtrap analysis was performed using random seed over multiple processors using function mcapply. Range of CI 0.8-0.99 and pAUC 0.8-0.95 was used to identify pathways associated with the PASC group. These pathways were used to differentiate the uninfected and PASC donors into separate clusters incorporating >50% of cluster size. The clustering was performed by the k-means approach implemented in ComplexHeatmap (v2.4) and visualized. The bootstrap analysis resulted in CI of 0.99 and pAUC of 0.95 which can differentiate uninfected and PASC donors in clusters. These parameters were used to identify pathways associated with PASC with a bootstrap of 1000 as mentioned before. The analysis resulted in 85 pathways. These 85 pathways then collapsed into 54 modules.

A module is defined if pairwise genests had an overlap of at least 25% (jaccard index 0.25) genes between them (Bader et al., 2010). The 54 modules then used to perform module enrichment at single sample level using GSVA. The normalized enrichment score for each module was scaled and clustered using K-means clustering implemented in ComplexHeatmap (v2.4) with parameter row_km and column_km. The identified clusters are then visualized in heatmap.

### Pathway enrichment analysis

Gene Set Enrichment Analysis (GSEA) ^24^ was performed among genes that defined early acute infection status and genes that defined longitudinal changes. A custom collection of genesets that included the Hallmark v7.2 genesets, KEGG v7.2 and Reactomev7.2 from the Molecular Signatures Database (MSigDB, v4.0) was used as the pathway database. The “Type III interferon signaling” gene set was manually curated from the Interferome database ^25^. Genes were pre-ranked by the decreasing order of their log fold changes or coefficients. The running sum statistics and Normalized Enrichment Scores (NES) were calculated for each comparison. The pathway enrichment p-values were adjusted using the Benjamini-Hochberg method and pathways with p-values < 0.05 were considered significantly enriched.

### Sample-level enrichment (SLEA)

Sample-level enrichment analysis ^26^ was used to represent the GSEA pathway expression results on a per-sample basis. The SLEA score was calculated by first calculating the mean expression value of proteins (averaged across single cells) enriched in a pathway, then comparing it to the mean expression of random sets of genes (averaged across single cells) of the same size for 1,000 permutations per sample. The difference between the observed and expected mean expression values for each pathway was determined as the SLEA pathway score per sample.

### Statistical analysis

All statistical analyses were performed using the corresponding functions in RStudio (version 4.1). Comparisons of single protein olink NPX or module ssGSEA scores between groups were tested using the Wilcoxon rank sum test and when appropriate, the Benjamini-Hochberg method was applied to adjust p-values in multiple-testing correction. Unless specified, an adjusted p-value of 0.05 was considered significant.

### Analysis of Su Y et al (2022) INCOV Olink data

The Olink proteomic data consisted of 204 SARS-CoV-2 (INCOV) patients and 289 healthy controls. The INCOV patients were studied at clinical diagnosis (T1), acute disease (acute, T2), and 2–3 months post onset of initial symptoms (convalescent, T3). Olink plasma proteomic data was available for a total of 443 proteins. Among these, 163 proteins overlapped with the differentially expressed proteins found in inflammatory signatures that significantly defined clusters 4 & 5 in our cohort. K-means unsupervised clustering of the INCOV Olink proteomic data was performed on the 163 protein overlap. To remain consistent with our cohort, we used samples available at the first timepoint ≥60 days PSO per INCOV patient (which made a total 74 INCOV patients). The kmeans function of the stats R package was used with k=5, allowing 100 iterations.

## ACKNOWLEDGEMENTS

We thank the study participants for their dedication to this project; the Allen Institute founder, Paul G. Allen, for his vision, encouragement, and support; Adam Savage and Tao Peng for review and helpful discussions of the final manuscript draft; Leila Shiraiwa and Nina Kondza for laboratory operations support, the Human Immune System Explorer (HISE) software development team at the Allen Institute for Immunology for their support and dedication. This paper and the research behind it would not have been possible without the collaborative computational data analysis environment provided by HISE.

## SUPPLEMENTARY MATERIALS

**Figure S1**. (A) Table of study participant demographics at the time of enrollment and the number and proportion of infected PASC patients with symptoms at >= 60 days PSO. (B) Longitudinal sampling timeline across infected patients. Serum was collected at 2-5 timepoints for each participant ranging from 6 to 379 days PSO. Patients are arranged by sex and increasing age for recovered and PASC.

**Figure S2**. (A) Hierarchical clustering heatmap of symptomatology across infected PASC patients. (B,C) Heatmap of the Olink serum proteins significantly associated with Fatigue/Malaise and pulmonary symptoms respectively among infected PASC patients. The uninfected and infected recovered patients were included to perform hierarchical clustering on these proteins. (D) Heatmap of the union of Olink serum proteins significantly associated with each of the symptoms among infected PASC patients. The uninfected and infected recovered patients were included to perform hierarchical clustering on these proteins.

**Figure S3**. (A) Stacked bar plot of the proportion of uninfected, recovered and PASC patients per cluster identified with the rule-in approach (B) (C,D) Box and jitter plots of SARS-CoV-2 specific CD4+ and CD8+ T-cells between the inflammatory PASC (PASC from clusters 4 and 5) and non-inflammatory PASC (PASC from clusters 2 and 3) respectively.

**Figure S4**. Modules that are significantly higher expressed in clusters 4 and 5 relative to all other clusters. Modules unique to a cluster are arranged and ranked by increasing adjusted p-value of <0.05, while modules expressed in both clusters are arranged and ranked by the average of their adjusted p-values. The color gradient of each node represents the -log10 adjusted p-value. P-values determined by Wilcoxon rank sum test are noted in supplementary table S5.

**Figure S5**. Box and jitter plots of the ssGSEA scores (y-axis) across all clusters (x-axis) for the modules that were significantly associated with each cluster. P-values determined by Wilcoxon rank sum test are noted in supplementary table S5.

**Figure S6**. Top 20 proteins ranked by adjusted p-value < 0.05 that are significantly higher expressed in cluster 4 (left) and cluster 5 (right) relative to all other clusters. The color gradient of each node represents the -log10 adjusted p-value. P-values determined by Wilcoxon rank sum test are noted in supplementary table S6.

**Figure S7**. Box and jitter plots of NPX (y-axis) across all clusters (x-axis) for the proteins that were significantly associated with each cluster. P-values determined by Wilcoxon rank sum test are noted in supplementary table S6.

**Figure S8**. Longitudinal protein expression (NPX as y-axis) of IFN-γ, TNF, its related cytokines and chemokines and type I interferon associated proteins on samples available from 6 days PSO through >60 days PSO (x-axis). PASC patients from the inflammatory clusters 4 and 5 are represented here as inflammatory PASC (red), PASC patients from clusters 2 and 3 are represented here as non-inflammatory PASC (blue) while the recovered patients are represented in black.

**Figure S9**. Longitudinal module expression (ssGSEA score as y-axis) of IFN-γ signaling, TNF signaling, its related modules and IFNa signaling on samples available from 6 days PSO through >60 days PSO (x-axis). PASC patients from the inflammatory clusters 4 and 5 are represented here as inflammatory PASC (red), PASC patients from clusters 2 and 3 are represented here as non-inflammatory PASC (blue) while the recovered patients are represented in black.

**Figure S10**. Box and jitter plots of the proteins significantly upregulated in the INCOV E vs INCOV from clusters B,C,D. P-values were determined by a Wilcoxon rank sum test. P-values determined by Wilcoxon rank sum test are noted in supplementary table S7.

## Author contributions

Conceptualization: TRT, XL, GLS, PJS, TFB, MJM

Methodology: AT, SVV, GLS, PJS, XL, TRT, JLC, MPL, ZM, KWC, HM, EWN, LBF, SCDR

Investigation: ZJT, PJS, AT, SVV, JLC, MPL, KWC, LBF, ZM, HM, XL, TRT

Visualization: AT, SVV, ZM

Funding acquisition: TFB, MJM

Project administration: TFB, MJM, LAB, EMC

Supervision: TRT, XL, PJS, EMC, TFB, MJM

Writing – original draft: TRT, AT

Writing – review & editing: TRT, AT, SVV, GLS, XL, TFB, MPL JLC, MJM

## Data and materials availability

All data are available in the main text or the supplementary materials. All data, code, and materials used in the analysis are available here: https://doi.org/10.5281/zenodo.6499388

