## Supplemental Figure 1 for "Persistent serum protein signatures define an inflammatory subset of long COVID"

A

| Group | Uninfected | Infected Recovered | Infected PASC |
|---|---|---|---|
| No. of participants | 22 | 24 | 55 |
| **Demographics** | | | |
| Sex - # (%) | | | |
| Female | 10 (45.5) | 15 (62.5) | 34 (61.8) |
| Male | 12 (54.5) | 9 (37.5) | 21 (38.2) |
| Age - median yr (range) | | | |
| Female | 44 (29-61) | 46 (20-79) | 52 (22-74) |
| Male | 52 (31-77) | 56 (24-65) | 47 (31-82) |
| **Symptoms >=60 Days PSO - #Yes (%), #No (%)** | | | |
| Fever and Chills | | | 3 (5.5), 52 (94.5) |
| Loss of Appetite or Weight Loss | | | 2 (3.6), 53 (96.4) |
| Fatigue/Malaise | | | 25 (45.5), 30 (54.5) |
| Pulmonary | | | 23 (41.8), 32 (58.2) |
| Cardiovascular | | | 8 (14.5), 47 (85.5) |
| Gastrointestinal | | | 3 (5.5), 52 (94.5) |
| Musculoskeletal | | | 9 (16.4), 46 (83.6) |
| Neurologic | | | 15 (27.3), 40 (72.7) |
| Tinnitus | | | 2 (3.6), 53 (96.4) |
| Any mild symptom | | | 35 (63.6), 20 (36.4) |

B

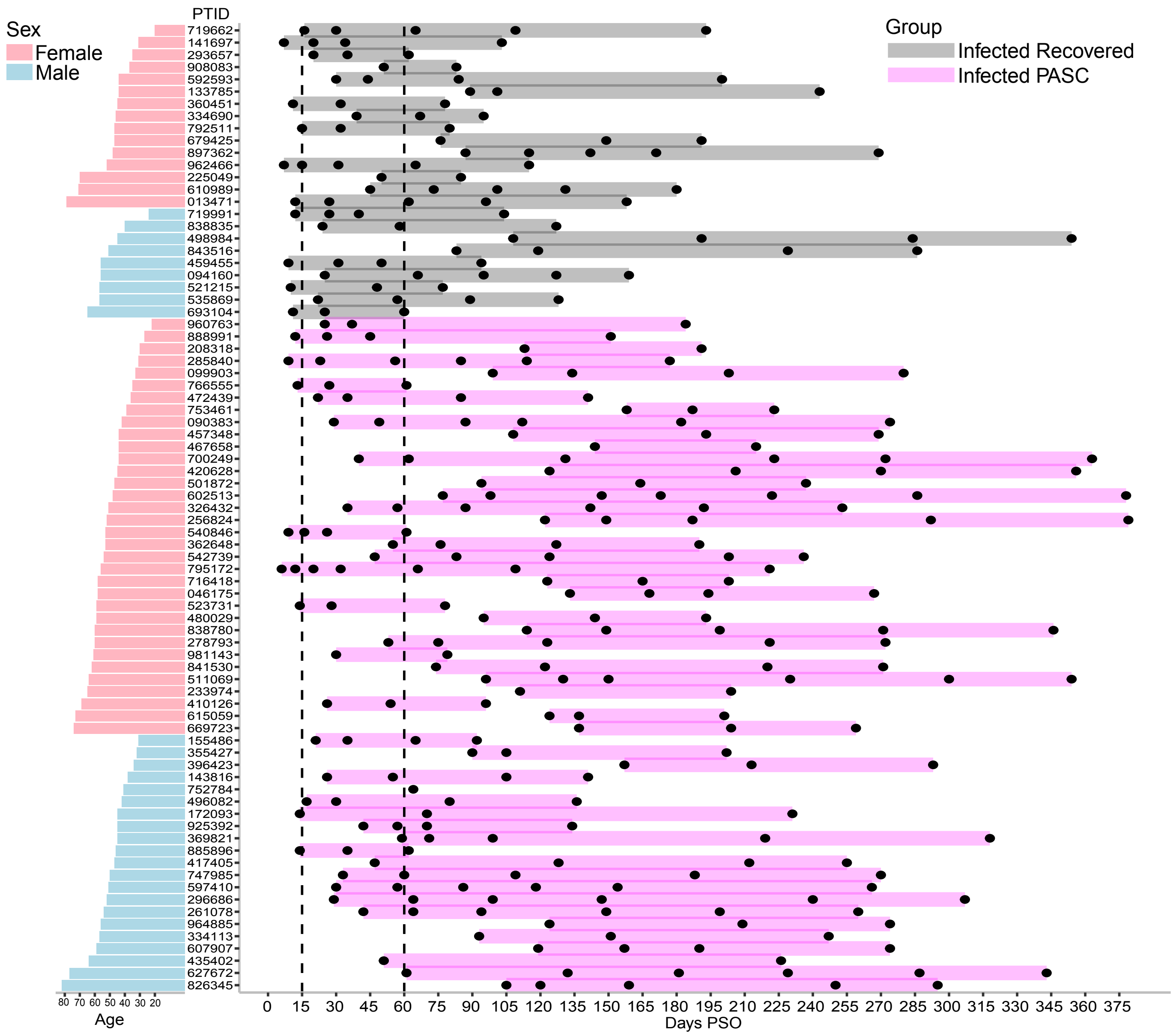
